# Distance-dependent reconfiguration of hubs in Alzheimer’s disease: a cross-tissue functional network study

**DOI:** 10.1101/2023.03.24.532772

**Authors:** Xingxing Zhang, Yingjia Li, Qing Guan, Debo Dong, Jianfeng Zhang, Xianghong Meng, Fuyong Chen, Yuejia Luo, Haobo Zhang, Alzheimer’s Disease Neuroimaging Initiative

## Abstract

The hubs of the intra-grey matter (GM) network were sensitive to anatomical distance and susceptible to neuropathological damage. However, few studies examined the hubs of cross-tissue distance-dependent networks and their changes in Alzheimer’s disease (AD). Using resting-state fMRI data of 30 AD patients and 37 normal older adults (NC), we constructed the cross-tissue networks based on functional connectivity (FC) between GM and white matter (WM) voxels. In the full-ranged and distance-dependent networks (characterized by gradually increased Euclidean distances between GM and WM voxels), their hubs were identified with weight degree metrics (frWD and ddWD). We compared these WD metrics between AD and NC; using the resultant abnormal WDs as the seeds, we performed seed-based FC analysis. With increasing distance, the GM hubs of distance-dependent networks moved from the medial to lateral cortices, and the WM hubs spread from the projection fibers to longitudinal fascicles. Abnormal ddWD metrics in AD were primarily located in the hubs of distance-dependent networks around 20-100mm. Decreased ddWDs were located in the left corona radiation (CR), which had decreased FCs with the executive network’s GM regions in AD. Increased ddWDs were located in the posterior thalamic radiation (PTR) and the temporal-parietal-occipital junction (TPO), and their FCs were larger in AD. Increased ddWDs were shown in the sagittal striatum, which had larger FCs with the salience network’s GM regions in AD. The reconfiguration of cross-tissue distance-dependent networks possibly reflected the disruption in the neural circuit of executive function and the compensatory changes in the neural circuits of visuospatial and social-emotional functions in AD.

## 1. Introduction

Evidence indicated that the blood oxygen level-dependent (BOLD) signals detected in the white matter (WM) were originated from neuronal activities in the grey matter (GM), which bring a series of biochemical and metabolic changes, including the exchanges of potassium and calcium ions and the production of nitric oxide, to evoke vascular hemodynamic responses in WM (1–4). A few studies examined the temporal synchronization of BOLD signals between WM bundles and GM regions, and found that cross-tissue functional connectivity (FC) was correlated with cognitive tasks performance (5–7). Recently, the cross-tissue FC has been applied in the studies of various diseases, including Alzheimer’s disease (AD) (8–11). One study found decreased cross-tissue FCs between the WM tracts and GM regions in AD patients, and their neuropsychological performance were significantly correlated with the FCs (12). The cross-tissue FCs between GM and WM were also used as classification features to increase the sensitivity and accuracy to differentiate AD from normal controls (NC) (11).

Abundant studies investigated the locations of the hubs in the intra-GM network, using weighted degree (WD) - a measure of FCs’ strength between one region and other regions, to define the hubs (13–15). Evidence showed that the hubs of intra-GM network were particularly vulnerable to neuropathological damage of AD (16, 17). Moreover, evidence suggested that the hubs were structurally sensitive, as the strength of FC was dependent on the distance between regions (7, 18–21). The short-ranged FCs and long-ranged FCs differed in the metabolic costs and susceptibility to neurodegenerative diseases, with the long-ranged FCs possibly consuming more resources and more prone to Aβ deposition (22, 23).

Despite the emergent applications of cross-tissue functional connections, the cross-tissue distance-dependent network’s topology was still less understood. Given the importance of the distance-dependent hubs, there was lack of a study on their configuration in cross-tissue distance-dependent networks in older adult and their changes in AD. Our study performed three levels of analysis to shed light on these issues. We constructed the cross-tissue networks based on the functional connections between GM and WM, and demonstrated the distribution patterns of two types of hubs (identified by high WD values) located in either GM or WM. Notably, the cross-tissue networks included the full-ranged ones and a series of distance-dependent ones characterized by gradually increased Euclidean distances between GM and WM voxels. Consequently, the WD metrics of full-ranged and distance-dependent networks were compared between AD and NC to reveal the regions with abnormally high or low WD in AD patients. Finally, to understand the roles of those regions in the pathological mechanism of AD, two further examinations were performed to illustrate how those abnormal WD metrics were composed by cross-tissue FCs and correlated with cognitive performance.

## 2. Materials and Methods

### 2.1 ADNI database

The present study used the open database, the Alzheimer’s Disease Neuroimaging Initiative (24), to acquire data. ADNI was launched in 2004 to investigate AD and its prodromal stages (http://www.adniinfo.org). ADNI included four databases (ADNI-1, ADNI-2, ADNI-3, ADNI-GO). An initial five-year study, termed ADNI-1, was followed by two renewal five-year studies termed ADNI-2 and −3; and ADNI-GO enrolled early MCI participants (25). As the ADNI-1 phase did not collect resting-state fMRI data and the ADNI-3 phase included the longitudinal data of ADNI-2, we only used the data from ADNI-2, consistent with previous studies on the database selection for resting-state fMRI data of AD patients (26, 27).

According to the standardized protocol, the ADNI data was collected from various acquisition sites across Canada, and the United States, approved by the Institutional Review Board at each acquisition site, and written informed consent was obtained from each participant.

### 2.2 Participants

The ADNI-2 imaging data were collected at multi-sites, with different scanners and parameters (24). To alleviate the scanner effect, we carefully checked the scanner information and only used the image data acquired by a 3.0T Philips scanner with the same acquisition parameters (see Section 2.3 for the parameters). The inclusion criteria for the participants: 1) a clinician-confirmed diagnosis of AD or ‘‘normal’’ at the screening visit; and 2) the complete resting-state fMRI data available for the participants at their first scan time in the ADNI 2 database. In total, 30 AD patients and 37 NC were included in this study. The sample sizes of AD and NC were comparable with previous AD studies that used ADNI resting-state fMRI data (26, 27). No significant differences were found between the two groups on age and gender (Table 1).

**Table 1.**
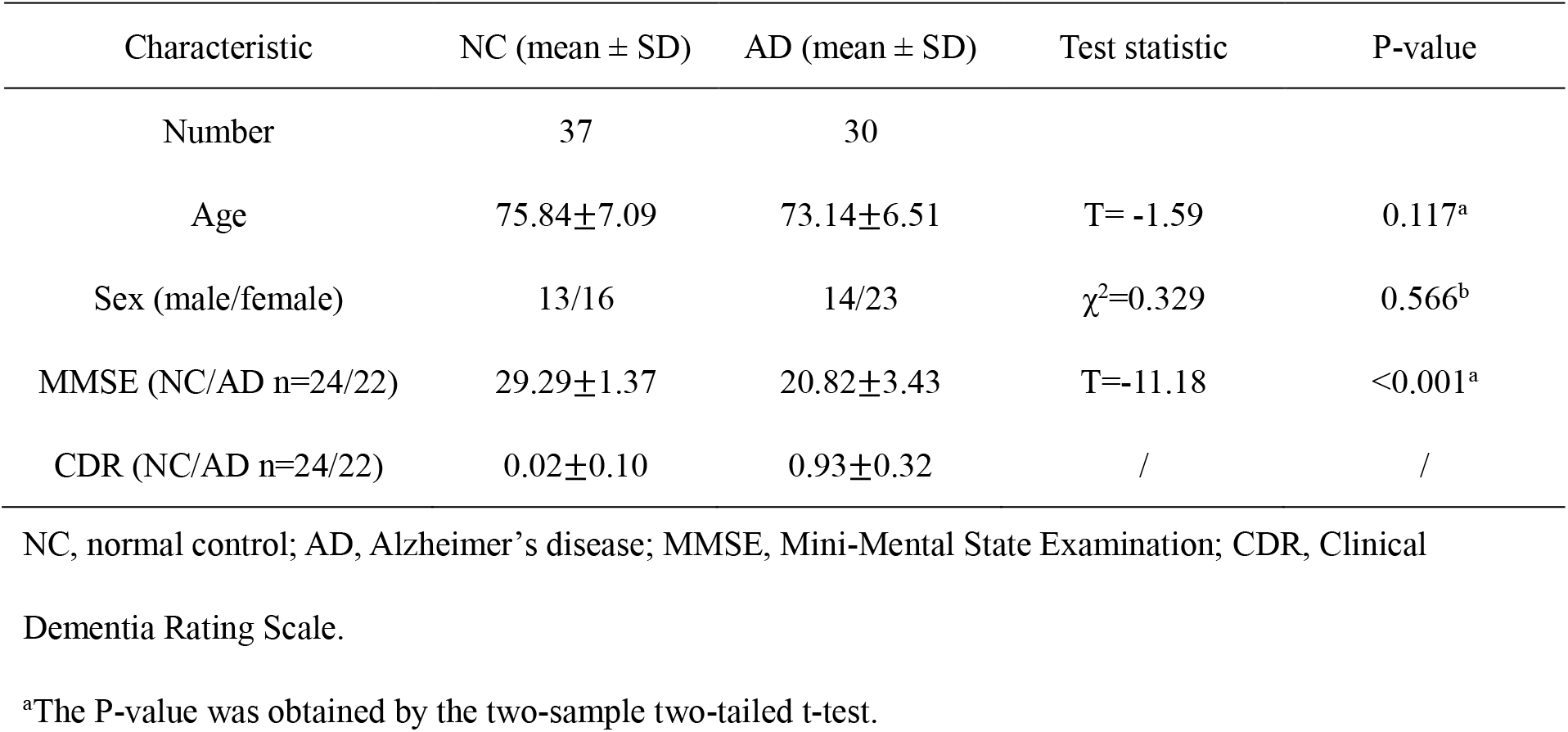
Demographic characteristics and neuropsychological tests.

### 2.3 Image acquisition and preprocessing

The resting-state fMRI data were obtained using an echo-planar imaging (EPI) sequence with the following parameters: repetition time (TR) = 3000 ms, echo time (TE) = 30 ms, flip angle = 80°, number of slices = 48, slice thickness = 3.313 mm, voxel size = 3 × 3 × 3 mm^3^, voxel matrix = 64 × 64, and total volume = 140. The slice order sequence was the same for all participants, as 1:2:47 and 2:2:48. The T1-weighted images were acquired using the following parameters: TR = 6.8 ms, TE = 3.1 ms, FA = 9°, slice thickness = 1.2 mm, number of slices = 170, voxel size = 1.1× 1.1×1.2mm^3^, acquisition matrix = 244×244.

Rs-fMRI images were preprocessed using the Data Processing and Analysis of Brain Imaging software package (DPABI, http://rfmri.org/dpabi) and the SPM12. The preprocessing procedure consisted of eight steps: 1) discarding the first ten volumes; 2) slice-timing; 3) head motion correction (the maximum motion threshold ≤2mm or 2°); 4) normalization and resampled to 3 × 3 × 3 mm^3^; 5) detrend; 6) nuisance covariates regression (Friston 24 for head motion, global signal, and cerebrospinal fluid signal regression); 7) temporal scrubbing (the scan volume with the frame-wise displacement (FD) >1 was removed); and 8) band-pass filter (0.01 Hz–0.15 Hz) to reduce low-frequency drift and high-frequency physiological noise (28, 29). One AD participant was removed from this study due to the maximum head motion beyond the threshold. Notably, we used the EPI template (provided by SPM, in standardized MNI space) for normalization, which generated satisfactory results. We also tried the normalization with T1 images using the DARTEL method, which generated poor quality results with parts of brain distortion, missing, or spatial mismatch. Previous AD studies have also used the EPI template for rs-fMRI normalization, instead of the T1 image, possibly due to severe structural atrophy of AD patients (4, 30–32).

### 2.4 GM and WM hubs and their WD metrics (WD-Gw and WD-Wg)

There were two types of hubs for each cross-tissue network constructed with the FCs between GM and WM regions. A GM hub was the GM region with strong FC connections to all WM regions; we used the metric of WD-Gw to identify them (i.e., the **W**eighted **D**egree of a given **G**M region by connecting to all **W**M regions). Vice versa, a WM hub would be the WM region with a high WD by connecting to all GM regions, identified by WD-Wg.

We used a voxel-wise approach to parcellate the brain as the neural signals extracted from the fine-grained units were more homogeneous and prone to the small-world property (33, 34). The masks of GM and WM tissues were generated with the two tissue probability maps provided by SPM12, respectively. The voxels with a GM probability-value >0.2 were defined as belonging to the GM mask (35) and a WM probability-value >0.6 as belonging to the WM mask (28, 29). Notably, 4405 voxels with a GM probability-value >0.2 & WM probability-value >0.6 were located in the boundary of two tissue masks. Given the ambiguous tissue type of these voxels, we excluded them from both masks. The locations of these overlapped voxels were shown in the supplementary materials (Fig s1).

### 2.5 Full-ranged and distance-dependent networks

#### 2.5.1 Full-ranged network and WD metrics (frWD-Gw & frWD-Wg)

We constructed the full-ranged cross-tissue networks to obtain the two WDs (i.e., frWD-Gw and frWD-Wg). The calculation pipeline is demonstrated in Fig 1A and Fig 1C. For voxel-based frWD-Gw, the Pearson’s correlation coefficients (Rs, with the threshold of R>0.2) of any given voxel in the GM mask with all voxels in the WM mask were calculated, Fisher r-to-z transformed and then summed. Similarly, for voxel-based frWD-Wg, the Rs (R>0.2) of any given voxel in the WM mask with all voxels in the GM mask were calculated, Fisher r-to-z transformed, and then summed. The whole-brain frWD-Gw/Wg maps of each individual were then smoothed (4mm) and z-standardized for the following statistical analysis.

**Fig 1.**
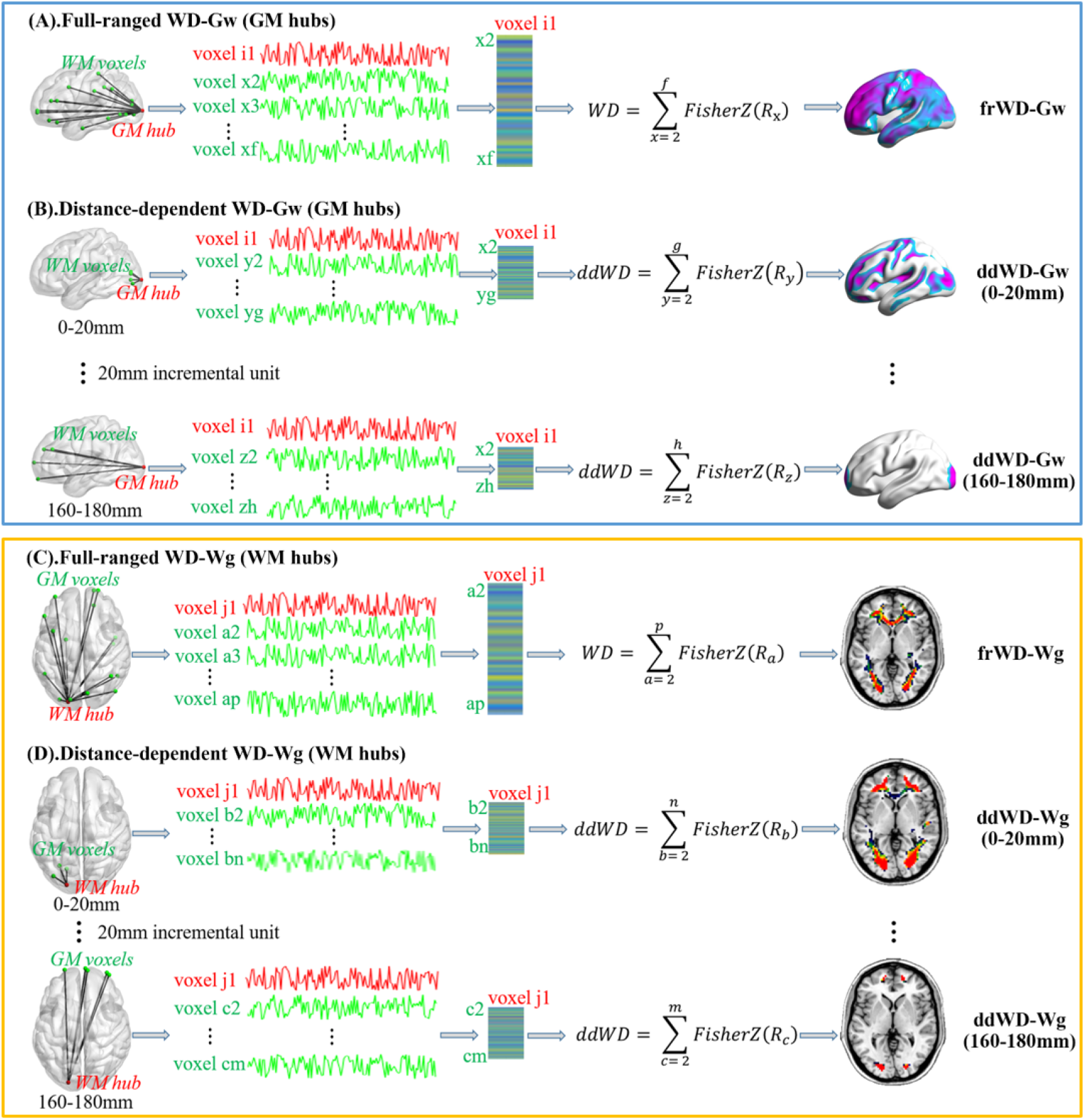
The calculation pipeline for weighted degree metrics in cross-tissue networks. (A) and (B) presented the calculation process to obtain the weighted degree of any given GM voxel by connecting to all WM voxels in full-ranged networks (frWD-Gw) and in distance-dependent networks based on the Euclidean distance between GM and WM voxels (ddWD-Gw); the GM hubs in these networks were identified with high WD-Gw values. Similarly, (C) and (D) showed the calculation process to obtain the weighted degree of any given WM voxel by connecting to all GM voxels in full-ranged networks (frWD-Wg) and in distance-dependent networks (ddWD-Wg); the WM hubs were identified with high WD-Wg values.

#### 2.5.2 Distance-dependent network and WD metrics (ddWD-Gw & ddWD-Wg)

Previous studies divided the Euclidean distance between any two voxels as short- and long-ranged with an arbitrary cut-off of 75mm (36, 37), or defined it with an incremental distance of 10mm (38). In this study, we used an incremental unit of 20mm to define the Euclidean distance of voxels into nine ranges, starting from 0 to 180mm (e.g., 0-20mm, 20-40mm… 160-180mm).

Consistent with previous studies on measuring the distance-dependent functional connectivity (38), the ddWD-Gw at each distance range was calculated as the sum of Fisher z-transformed Rs (R>0.2) of any given GM voxel connecting to the WM voxels whose Euclidean distances to that specific GM voxel were constrained within that range. Similarly, the ddWD-Wg at each distance range was calculated as the sum of Fisher z-transformed Rs (R>0.2) of any given WM voxel connecting to the GM voxels whose Euclidean distances to that specific WM voxel were constrained within that range. These ddWD maps were also smoothed (4mm) and z-standardized for statistical analysis. The calculation pipeline for ddWD metrics is demonstrated in Fig 1B and Fig 1D.

### 2.6 Seed-based cross-tissue FC

To illustrate the cross-tissue FCs’ composition of abnormal WD metrics in AD patients, we performed seed-based cross-tissue FC analysis. Firstly, we defined each supra-threshold cluster from the results of comparison analysis on different WD metrics as the seed. Secondly, the average time series of voxel-based BOLD signals within each seed were calculated from the pre-processed rs-fMRI images using the DPABI software (Data Processing & Analysis for Brain Imaging, *http://rfmri.org/dpabi*) (39). Thirdly, the seed’s average time series were correlated with the time series of the *targeting* voxels located in the tissue mask different from the seed. For example, if the seed was located in the GM, the *targeting* voxels would be in the WM mask, and vice versa. Notably, when the seed was derived from the analysis of ddWD at a certain range, the seed-based distance-dependent functional connectivity (ddFC) would be obtained between the seed and the *targeting* voxels whose Euclidean distances to any voxels of the seed were constrained within that range. The seed-based FCs also underwent the Fisher r-to-z transformation and smoothed (4mm).

### 2.7 Additional analysis: intra-tissue network

To have a comprehensive understanding of the topology of different functional networks in older adults and AD patients, we also established two intra-tissue networks within GM and WM, respectively, and obtained their full-ranged (frWD-G and frWD-W) and distant-dependent WD metrics (ddWD-G and ddWD-W). The calculation pipeline for intra-tissue network and their WDs metrics were described in the supplementary materials.

### 2.8 Statistical analysis

Our first analysis was to demonstrate the locations of cross-tissue network’s hubs in NC and AD groups. We obtained the *mean network* of each group by averaging the WD values across all individuals’ networks. The hubs in the *mean network* were defined as the voxels with z-standardized WD value ≥1 (calculated by subtracting the mean WD and then divided by the standard deviation of WDs of all voxels within the network), based on the previously used method of defining the hubs in functional GM networks (15, 23).

Subsequently, we compared AD with NC on different WD metrics (including frWD-Gw and frWD-Wg, ddWD-Gw and ddWD-Wg at different ranges), implemented on SPM12. Based on the resultant supra-threshold clusters, we then obtained the seed-based cross-tissue FCs and compared them between AD and NC. The controlled covariates for every group comparison analysis were the same, including age, gender, and mean FD of head motion. The significance threshold was set at a voxel-level threshold of p<0.005 (uncorrected) combined with a cluster-level threshold of p<0.05 (FWE-corrected), based on the Gaussian Random Fields theory (GRF) to correct for multiple comparisons. Notably, when the seed-based FC analysis did not show significant results under the above threshold, we lowered the significance threshold to a voxel-level p<0.05 (uncorrected) combined with a cluster size of 40 voxels. To validate the cognitive relevance of the regions with abnormal WD metrics, we performed the correlation analysis between MMSE and the mean WD metrics of the seeds controlled for age, gender, and mean FD of head motion, implemented on SPSS 20.0. The results are shown in the scatter plot (Fig 4).

**Fig. 2.**
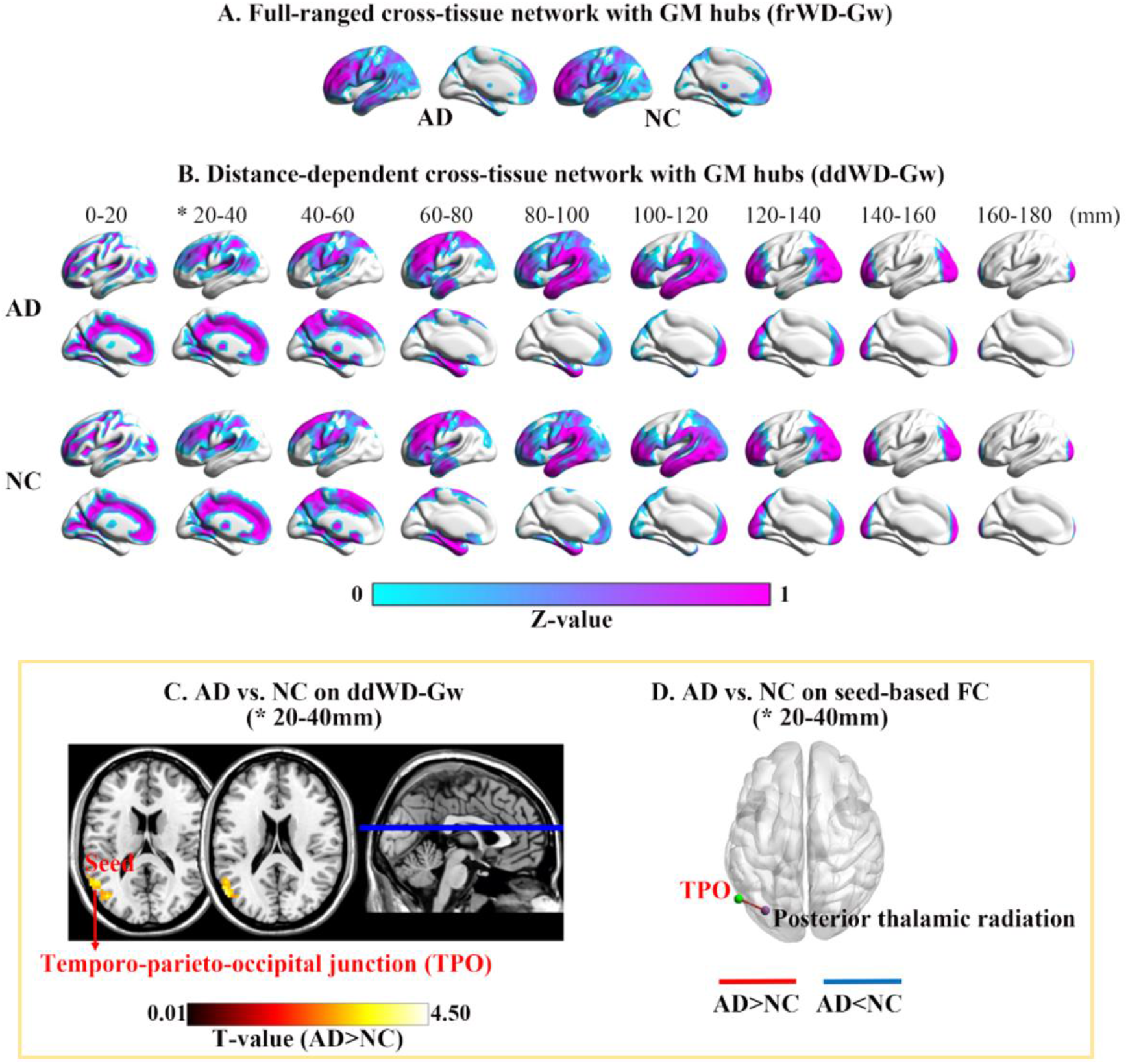
Mean maps of WD-Gw metrics in two groups and their difference between groups. * indicated significant differences between AD patients and NC in WD-Gw metrics. Maps (A) and (B) showed the mean WD-Gw maps of each group in full-ranged networks and distance-dependent networks, respectively. The GM hubs in each mean map were defined as the GM voxels with z-standardized WD value ⩾1. (C) Voxel-based WD-Gw metrics were compared between AD and NC, and the significantly increased ddWD-Gw were found at 20-40mm in AD. (D) Using the supra-threshold cluster from (C) as the seed, the FCs between the seed (in green) and the targeting WM region (in purple) were larger in AD.

**Fig 3.**
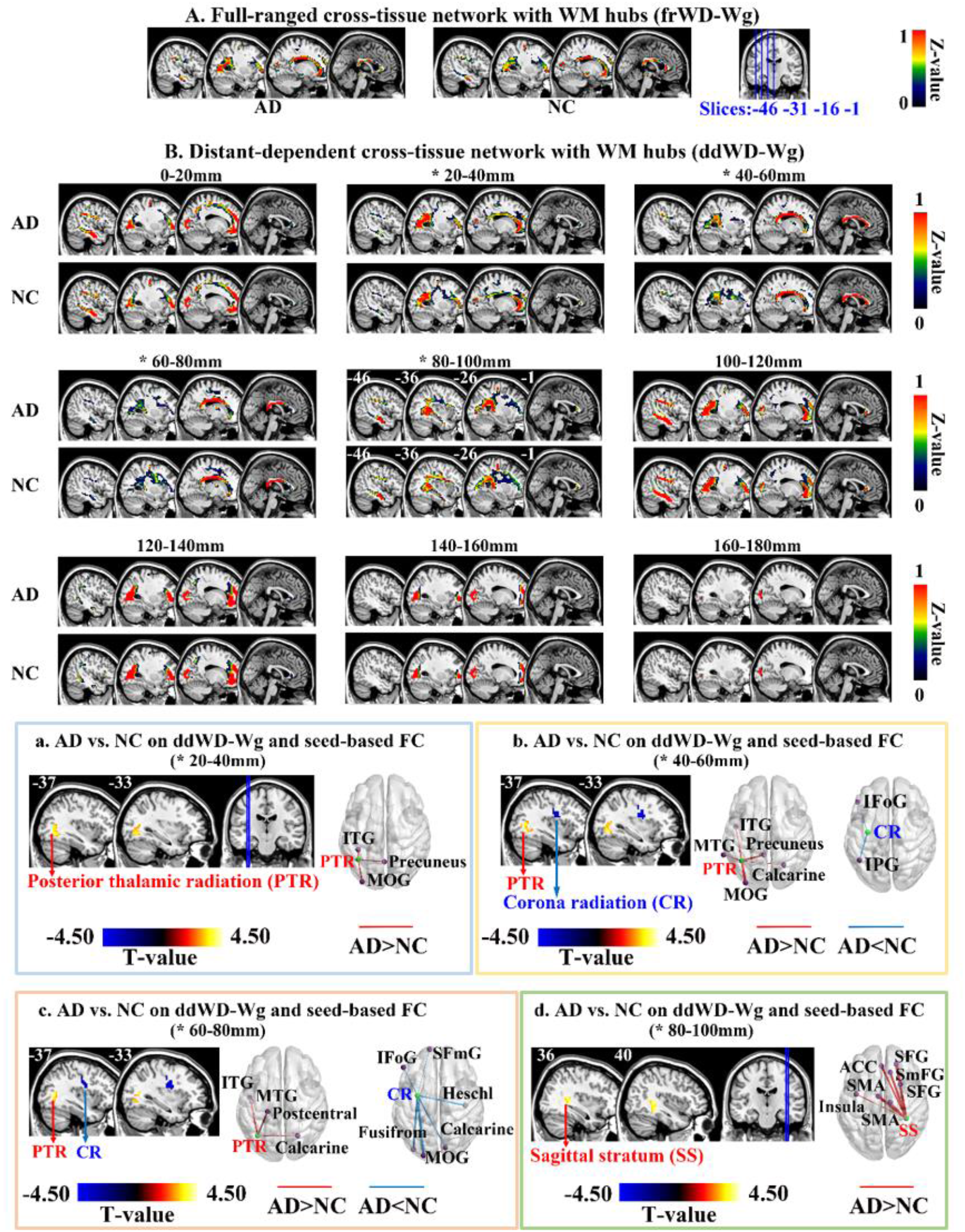
Mean maps of WD-Wg metrics in two groups and their difference between groups. * indicated significant differences between AD patients and NC in WD-Wg metrics. Maps (A) and (B) showed the mean WD-Wg maps of each group in full-ranged networks and distance-dependent networks, respectively. The WM hubs in each mean map were defined as the WM voxels with z-standardized WD value ⩾1. Each of Fig 3(a)-(d) contained the group comparison results on ddWD-Wg (sagittal view in the left) and seed-based FC metrics (3D view in the right). The FCs between the seed (in green) and targeting GM regions (in purple) were different between AD and NC.

**Fig 4.**
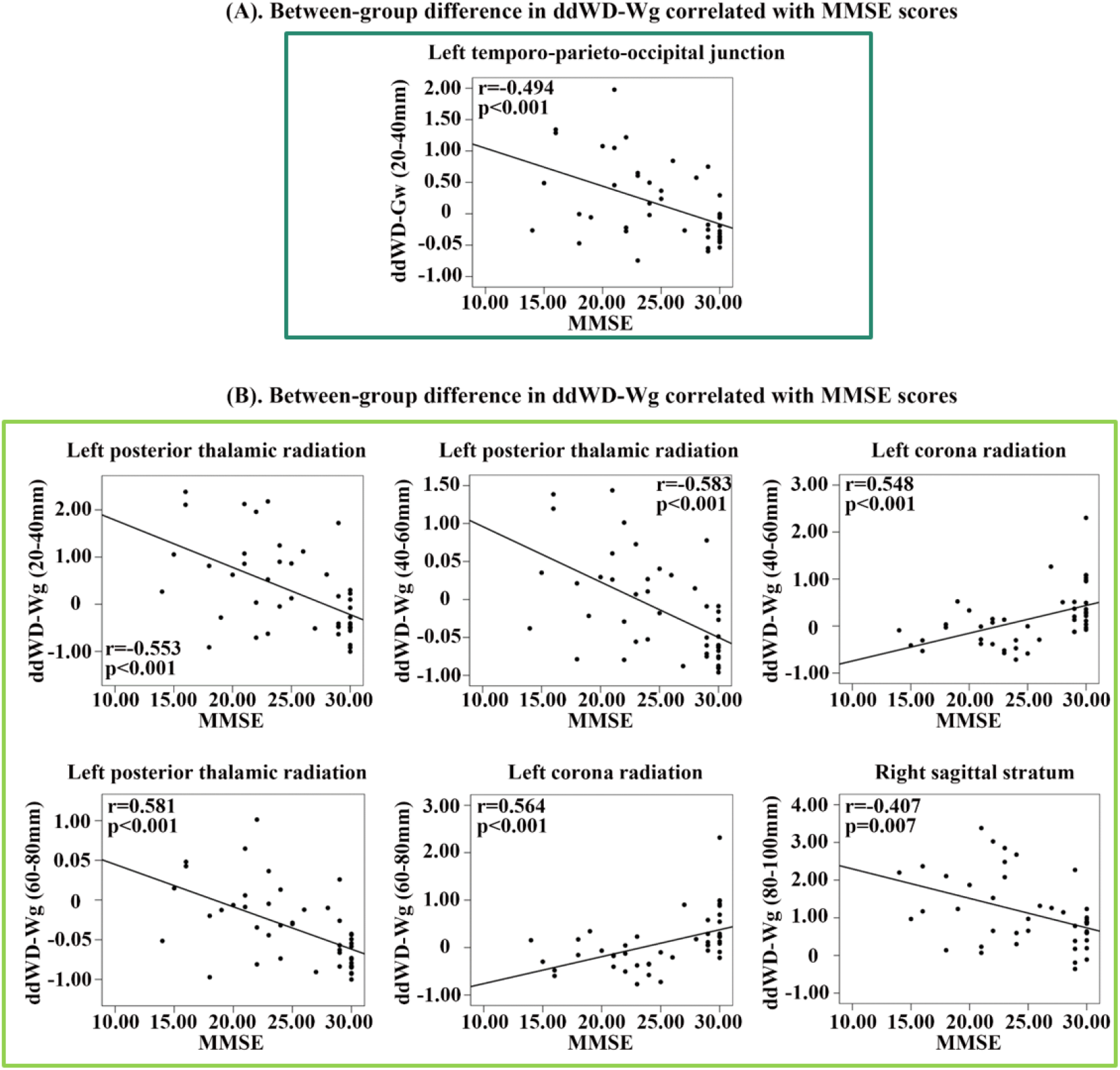
Correlation analysis between cross-tissue network metrics and MMSE scores. We performed the correlation analysis between MMSE and the mean WD metrics of the seeds, which were showed by the scatter plot. (A) the significant linear correlation between the ddWD-Gw and MMSE scores in all participants. (B) the significant linear correlation between the ddWD-Wg and MMSE scores in all participants in all participants.

Additionally, we demonstrated the hub distribution patterns in two intra-tissue networks and compared AD and NC on their WD metrics, using the same covariates and significance threshold as those in the cross-tissue network.

## 3. Results

### 3.1 Hub location of cross-tissue networks

For the cross-tissue full-ranged networks, the GM hubs (z-standardized frWD-Gw ⩾ 1) were mainly located in the lateral and medial frontal cortices, superior parietal and occipital cortices. For the distant-dependent networks, the spatial distributions of GM hubs (z-standardized ddWD-Gw ⩾ 1) changed along with incremental distances, moving gradually from the medial to lateral areas. The GM hubs of ddWD-Gw networks at the distance of 0-60mm were located in the bilateral medial cortices (medial prefrontal and cingulate cortices, precuneus, and calcarine), sulci, temporo-parieto-occipital (TPO) junction, and subcortical regions (caudate and thalamus). The GM hubs at the distance of 60-120mm were distributed in the lateral frontal, temporal and parietal cortices. The GM hubs at the distance of 120-180mm were restricted to the frontal and occipital poles (Fig 2).

For the cross-tissue full-ranged networks, the WM hubs (z-standardized frWD-Wg ⩾ 1) were mainly located in the projection fibers of posterior thalamic radiation (PTR) and corona radiation (CR), and commissural fibers (i.e., corpus callosum). For the distant-dependent networks, the WM hubs (z-standardized ddWD-Wg⩾1) showed four distribution patterns along with the incremental distance ranges. For the ddWD-Wg networks at the distance of 0-40mm, the WM hubs were mainly located in the two projection fibers (PTR and CR). The WM hubs at the distance of 40-80mm were located in the corpus callosum and PTR. The WM hubs at the distance of 80-120mm were mainly located in the association fibers (sagittal striatum and longitudinal fasciculus). Finally, the WM hubs at the distance of 120~180mm were restricted to the PTR and CR (Fig 3).

### 3.2 WDs of cross-tissue networks between AD and NC

Compared on the full-ranged WD metrics (i.e., frWD-Gw and frWD-Wg), no significant differences were found between AD and NC.

On the distance-dependent WD metrics, AD patients showed increased ddWD-Gw at the distance of 20-40mm, located in the left TPO junction (Table 2 and Fig 2). AD patients showed decreased ddWD-Wg at 40-60mm and 60-80mm, both located in the left CR (Table 3 and Fig 3). AD patients also showed increased ddWD-Wg at three distances (20-40mm, 40-60mm, 60-80mm), all located in the left PTR, and increased ddWD-Wg at the distance of 80-100mm, located in the right sagittal stratum.

**Table 2.**
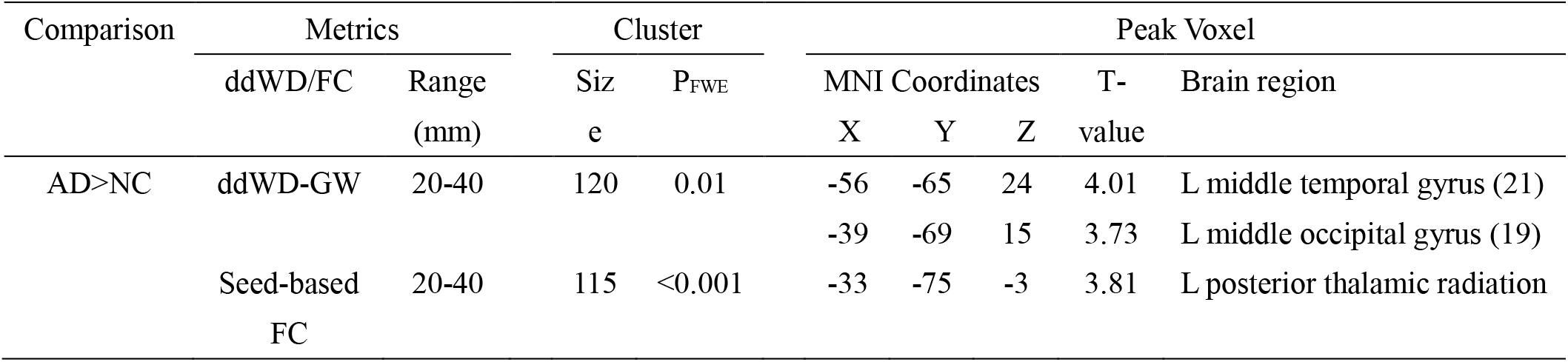
Differences between AD patients and NC on ddWD-GW and the seed-based FC. Voxel-based ddWD-GWs at nine ranges of Euclidean distance were compared between AD patients and NC. Using the resultant supra-threshold cluster as the seed, the seed-based FCs were compared between AD patients and NC. The adjusted covariates included age, gender, mean frame-wise displacement for head motion. The significance threshold was set at voxel-level p<0.005 (uncorrected) with a cluster-level p<0.05 (FWE-corrected).

**Table 3.** Difference between AD patients and NC on ddWD-WG and the seed-based FC. Voxel-based ddWD -W Gs at nine ranges of Euclidean distance were compared between AD patients and NC. Using the resultant supra-threshold clusters as the seed, the seed-based functional connectivity (FC) was compared between AD patients and NC. The adjusted covariates included age, gender, mean frame-wise displacement for head motion. The significance threshold was set at voxel-level p<0.005 (uncorrected) with a cluster-level p<0.05 (FWE-corrected).

|  | Metrics |  | Cluster |  | Peak Voxel |  |  |  |  |
| --- | --- | --- | --- | --- | --- | --- | --- | --- | --- |
|  | ddWD/FC | Range<br>(mm) | Size | P<sub>FWE</sub> | MNI |  |  | T-value | Brain region (BA) |
|  |  |  |  |  | X | Y | Z |  |  |
| AD<NC | ddWD-WG | 40-60 | 55 | 0.034 | -33 | -12 | 21 | -4.13 | L corona radiation |
|  | ddWD-WG | 60-80 | 59 | 0.022 | -30 | -3 | 30 | -3.68 | L corona radiation |
| AD>NC | ddWD-WG | 20-40 | 64 | 0.022 | -39 | -57 | -3 | 3.73 | L posterior thalamic radiation |
|  | ddWD-WG | 40-60 | 75 | 0.008 | -39 | -57 | -3 | 3.66 | L posterior thalamic radiation |
|  | ddWD-WG | 60-80 | 49 | 0.047 | -36 | -63 | 9 | 3.94 | L posterior thalamic radiation |
|  | ddWD-WG | 80-100 | 55 | 0.029 | 36 | -39 | 3 | 3.87 | R sagittal stratum |
| AD>NC | Seed-based FC | 80-100 | 64 | 0.038 | -39 | -3 | 9 | 3.61 | L insula |
|  |  |  | 569 | <0.001 | 21 | 42 | 36 | 4.60 | R superior frontal gyrus (9) |
|  |  |  |  |  | 12 | 30 | 57 | 4.37 | R superior medial frontal gyrus (8) |
|  |  |  |  |  | -1 | 39 | 24 | 4.34 | L anterior cingulate cortex (32) |
|  |  |  | 76 | 0.018 | -6 | -6 | 60 | 4.45 | L supplementary motor area (6) |
|  |  |  |  |  | 12 | -15 | 69 | 3.51 | R supplementary motor area (6) |
|  |  |  | 64 | 0.038 | 27 | 12 | 57 | 3.83 | R superior frontal gyrus (8) |

The correlation analysis with MMSE validated the comparison results between AD and NC, as MMSE was positively correlated with the WD metrics that were decreased in AD patients, and negatively correlated with the WD metrics that were increased in AD patients (Fig 4).

### 3.3 Seed-based cross-tissue FC between AD and NC

Using the resultant regions with abnormal ddWD changes as the seeds, we obtained the seed-based ddFCs and then compared them between two groups. For the seed in the left TPO junction with increased ddWD-Gw at 20-40mm in AD, the ddFCs that connected the seed to the left PTR were significantly larger in AD patients (Table 2 and Fig 2). For the seed in the right sagittal striatum with increased ddWD-Wg at 80-100mm in AD, the ddFCs that connected the seed to the following GM regions were significantly larger in AD patients, including the left insula, left anterior cingulate cortex, right superior frontal gyrus, right medial superior frontal gyrus, and bilateral supplementary motor area (Table 3 and Fig 3).

For five seeds, their seed-based ddFCs showed differences between AD and NC under a more lenient threshold (uncorrected voxel-level p<0.05 & cluster size ≥ 40 voxels). Of the two seeds in the left CR with decreased ddWD-Wg at 40-60mm and 60-80mm in AD, the ddFCs that connected the seeds to the left prefrontal cortices and inferior parietal gyrus were smaller in AD. Of the three seeds in the left PTR with increased ddWD-Wg at 20-40mm, 40-60mm, and 60-80mm in AD, the ddFCs that connected the seeds to the left inferior temporal gyrus, precuneus, and occipital cortices were larger in AD (Supplementary Table s3 and Fig 3).

### 3.4 WDs of intra-tissue networks between AD and NC

The distribution patterns of intra-GM and intra-WM networks’ hubs were demonstrated in the Supplementary Fig s2 and Fig s3. We compared AD and NC on the WD metrics of intra-tissue networks (including frWD-G and frWD-W, ddWD-G and ddWD-W); AD patients showed decreased ddWD-G at 0-20mm in the bilateral thalamus and decreased ddWD-W at 20-40mm in the left thalamus and subthalamic nucleus (Supplementary Table s1,Table s2 and Fig s4).

## 4. Discussion

### 4.1 Hub distribution of cross-tissue networks

In older adults (including NC and AD), the GM hubs of full-ranged cross-tissue network (i.e. frWD-Gw) were mainly located the dorsal lateral prefrontal cortices, superior and inferior parietal lobules, which were the central regions of the fronto-parietal network and dorsal attention network (40). The GM hubs of distance-dependent cross-tissue networks (ddWD-Gw) gradually moved from the deeper cortical areas to the surface with incremental distances. Such distribution patterns were similar to the hubs of distant-dependent intra-GM network, as shown in the study by Dai (13) and our study (see the supplementary Fig.s3). In Dai’s and our studies, the distance ranges were both short, 10mm and 20mm, respectively. Previous studies indicated that the BOLD signals detected in WM regions were possibly originally from their neighboring GM neuronal activities (7, 19). Hence, the FC of a GM region with another GM region or its adjacent WM region could be much similar, resulting in the observed resemblance between the GM hub’s distribution of the cross-tissue networks and that of the intra-GM networks.

The WM hubs of full-ranged cross-tissue network (i.e. frWD-Wg) were located in the two projection fibers (PTR and CR) and the commissural fibers of corpus callosum. In ddWD-Wg networks, with increasing distance, the distribution of hubs moved from the projection fibers, to the commissure fibers, then to the association fibers. Such distribution patterns of WM hubs might be consistent with their structural and functional roles in the brain. The PTR and CR contained the projection fibers connecting the subcortical regions to their neighboring cortical regions, and are involved in multiple cognitive processes (41). The middle-length fibers of corpus callosum connect the bilateral hemispheres to allow fast communication (42, 43). The sagittal stratum contains the various longitudinal fascicles, connecting the extensive areas across the frontal, temporal, parietal, and occipital lobes, and involved in attention, executive control, and affective processes (44–48).

### 4.2 Topological change of cross-tissue networks in AD patients

Abnormally increased and decreased ddWD metrics were found in AD patients, primarily located in the hub areas of cross-tissue networks, which included the TPO (GM hub) and the PTR, CR and sagittal striatum (WM hubs). Complemented by the results of seed-based FC analysis, we could demonstrate the specific regions whose cross-tissue FCs with the hubs contributing to AD-related changes in ddWDs. Based on these findings, the implications of these cross-tissue connections in the pathological mechanism of AD were discussed.

#### 4.2.1 Decreased ddWD-Wg in the CR

The present study showed that the decreased ddWD-Wg of AD patients was located in the left CR at 40-60 mm and 60-80mm; and the ddFCs that connected the left CR to the left prefrontal and inferior parietal cortices were also decreased in AD. The CR connected the striatum and thalamus with the frontal and parietal areas (49). Evidences indicated that the CR’s structural integrity was associated with processing speed and cognitive flexibility across life (49–52). The prefrontal and inferior parietal cortices are the key regions of executive network and involved in cognitive control (53–55). Previous studies showed that the structural integrity of CR was disrupted in AD, evidenced by decreased fractional anisotropy (56–58). Our results regarding the decreased ddWD-Wg in the left CR possibly reflected the damage in the neural circuit of cognitive control in AD patients.

#### 4.2.2 Increased ddWD-Gw in the TPO

Our results showed that AD patients had larger ddWD-Gw at 20-40mm in the left TPO junction than NC, and the ddFCs between this region and the left PTR were larger in AD patients than NC. As a hub region of ddWD-Gw network at 20-40mm, the TPO junction was located in a converging site of the temporal, parietal, and occipital cortices, and involved in several high-level cognitive functions, including visuospatial function, face and object recognition, and language (59). The PTR connected the caudal thalamus with the posterior parietal and occipital cortices (60–62), acting as the anatomical foundation of visuospatial function. A previous study showed that neural activity level in the TPO areas during a clock-drawing task was higher in the early stage of AD and declined in the late stage of AD (63). According to the MMSE score, AD could be divided into mild, moderate, and severe stages (MMSE: 19-24, 10-18, 0-9) (64). As such, our participants were mainly in the early stage of AD (MMSE: 20.82±3.43). Previous studies proposed a compensation mechanism to account for enhanced neural activity and increased FCs observed in older adults, suggesting that the plastic brain could reorganize functional circuits to offset the age-related neural inefficiency (65–68). Our results on the increased ddWD-Gw in the TPO in AD were consistent with these lines of evidence, supporting the compensation mechanism to offset the decline of visuospatial function in early AD (60, 63, 69, 70).

#### 4.2.3 Increased ddWD-Wg in the PTR

We also found increased ddWD-Wg at 20-40mm, 40-60mm, and 60-80mm in the left PTR in AD; the seed-based FC analysis revealed increased ddFCs that connected the left PTR and the left inferior temporal gyrus, precuneus, and occipital cortices in AD. Enhanced FCs in the occipital cortices in AD patients were also observed in Dai’s study that compared the intra-GM network metrics between AD and NC (13). As explained in the above section, the PTR was essential in visuospatial function by connecting the caudal thalamus with the parietal and occipital cortices (71). The inferior temporal gyrus and occipital cortices played important roles in visuospatial function. The precuneus was also involved in visuospatial processing and episodic memory retrieval (72). The increased ddWD-Wg in the PTR in AD patients might indicate the compensated changes in the visuospatial neural circuits in our AD participants.

#### 4.2.4 Increased ddWD-Wg in the sagittal stratum

In our study, AD patients showed larger ddWD-Wg at 80-100mm in the right sagittal stratum; increased ddFC were found between this WM region and the GM regions of insula, anterior cingulate, supplementary motor, and medial frontal cortices, which were all the core regions of the salience network (73, 74). The sagittal stratum contains the middle and inferior longitudinal fascicles and inferior fronto-occipital fascicle (75). By connecting with the long-ranged regions across the frontal, temporal, parietal, and occipital lobes, the sagittal stratum has been involved in multiple mental processes, including the affective process (44, 46–48). Through the connection with the amygdala (76), the salience network is activated in response to emotionally significant internal and external stimuli (77). Emotional deregulation, a common symptom in MCI and AD, occurred in 35-85% of MCI individuals and up to 75% of AD patients. Convergent evidence demonstrated that the hyper-connectivity of the salience network was associated with affective disorders in AD, such as anxiety, irritability, aggression, and euphoria (78, 79). Our finding identified the abnormal increase in the cross-tissue FCs between the sagittal stratum and the salience network, which might be the neural substrates for the affected social-emotional processing in AD.

### 4.3 Biomarkers from distant-dependent cross-tissue networks

Abundant functional network metrics have been employed as neuroimaging biomarkers in clinical applications. Intriguingly, we found only the metrics of distance-dependent cross-tissue networks (i.e., ddWD-Gw/Wg) at 20-100mm showed significant difference between AD and NC, while the metrics of full-ranged cross-tissue networks (i.e., frWD-Gw/Wg) or the metrics of intra-tissue networks (i.e. frWD-G/W and ddWD-G/W) showed none or little. The sensitivity of distance-dependent network metrics in capturing disease-related changes or cognitive relevance have been noted in previous studies (13, 14), indicating that the anatomical distance was a key factor to characterize FC. For example, the sagittal stratum contained the longitudinal fascicles that connected widespread cortical regions; hence, it would be easier to reveal the abnormal changes occurred in this region by using the long-ranged FC metrics. Regarding the difference in cross-tissue and intra-tissue networks, a recent study demonstrated that using the cross-tissue FC achieved a superior classification accuracy between AD and NC, compared to using the FCs of intra-GM network (11). Our findings suggested that cross-tissue metrics were more susceptible to AD-related changes, which could induce a network’s reconfiguration as evident in those reduced and enhanced functional connections (80, 81).

## 5. Limitation

The present study had a few limitations. Firstly, the sample size was relatively small, with 30 AD patients and 37 NC in the ADNI database eligible to our study. Secondly, the incomplete neuropsychological assessments for these participants restricted our correlation analyses to MMSE only. Thirdly, the ddWD metrics in this study reflected stable synchronization of cross-tissue BOLD signals across time. However, the characteristics of temporal synchronization could be dynamic between the two tissues, which would be of much interest for future exploration in AD patients.

## 6. Conclusion

The hubs of cross-tissue distance-dependent networks showed distinct distribution patterns in two tissues: the GM hubs moved from the medial (distance <60mm) to lateral cortices (distance >60mm), while the WM hubs spread from the projection fibers (distance <40mm), the commissure fibers (distance 40-80mm), then to the longitudinal fascicles (distance >40mm). The cross-tissue ddWD metrics were sensitive biomarkers to capture abnormal changes in AD, with decreased and increased ddWD metrics located in the hubs of cross-tissue networks around 20-100mm. Through the reconfiguration of cross-tissue networks, these hubs were possibly involved in the damage in executive function, and the compensatory changes in visuospatial and social-emotional functions of AD patients.

## 7. Author contributions

Xingxing Zhang: conceptualization, methodology, writing the original draft. Yingjia Li: draw maps, review & editing. Qing Guan: review & editing. Debo Dong: review & editing. Fuyong Chen: review & editing. Jing Yi: review & editing. Yuejia Luo: review & editing. Haobo Zhang: conceptualization, methodology, writing, review & editing.

## 8. Acknowledgements

This study was supported by the National Natural Science Foundation of China (No. 31700960, 32071100), the Natural Science Foundation of Guangdong Province of China (2020A1515011394), the Shenzhen Fundamental Research General Project (JCYJ20190808121415365), the Shenzhen-Hong Kong Institute of Brain Science-Shenzhen Fundamental Research Institutions grant (2019SHIBS0003), and the National Key Research and Development Program of China (2018YFC1315205). The funders had no role in study design, data collection and analysis, decision to publish, or preparation of the manuscript.

The data used in this study was from the Alzheimer’s Disease Neuroimaging Initiative (24) (National Institutes of Health Grant U01 AG024904) and DOD ADNI (Department of Defense award number W81XWH-12-2-0012). ADNI is funded by the National Institute on Aging, the National Institute of Biomedical Imaging and Bioengineering, and through generous contributions from the following: AbbVie, Alzheimer’s Association; Alzheimer’s Drug Discovery Foundation; Araclon Biotech; BioClinica, Inc.; Biogen; Bristol-Myers Squibb Company; CereSpir, Inc.; Cogstate; Eisai Inc.; Elan Pharmaceuticals, Inc.; Eli Lilly and Company; EuroImmun; F. Hoffmann-La Roche Ltd. and its affiliated company Genentech, Inc.; Fujirebio; GE Healthcare; IXICO Ltd.; Janssen Alzheimer Immunotherapy Research & Development, LLC.; Johnson & Johnson Pharmaceutical Research & Development LLC.; Lumosity; Lundbeck; Merck & Co., Inc.; Meso Scale Diagnostics, LLC.; NeuroRx Research; Neurotrack Technologies; Novartis Pharmaceuticals Corporation; Pfizer Inc.; Piramal Imaging; Servier; Takeda Pharmaceutical Company; and Transition Therapeutics. The Canadian Institutes of Health Research is providing funds to support ADNI clinical sites in Canada. Private sector contributions are facilitated by the Foundation for the National Institutes of Health (www.fnih.org). The grantee organization is the Northern California Institute for Research and Education, and the study is coordinated by the Alzheimer’s Therapeutic Research Institute at the University of Southern California. ADNI data are disseminated by the Laboratory for Neuro Imaging at the University of Southern California.

## 10. Data for reference

All the data was obtained from openly available data sets. Code will be deposited in an open access platform, upon acceptance of this manuscript.

## Reference

1. Gawryluk JR, Mazerolle EL, D’Arcy RCN. Does functional MRI detect activation in white matter? A review of emerging evidence, issues, and future directions. Frontiers in neuroscience. 2014;8.

2. Smith AJ, Blumenfeld H, Behar KL, Rothman DL, Shulman RG, Hyder F. Cerebral energetics and spiking frequency: the neurophysiological basis of fMRI. Proceedings of the National Academy of Sciences of the United States of America. 2002;99(16):10765–70.

3. Marcar VL, Loenneker T. The BOLD response: a new look at an old riddle. Neuroreport. 2004;15(13):1997–2000.

4. Calhoun VD, Wager TD, Krishnan A, Rosch KS, Seymour KE, Nebel MB, et al. The Impact of T1 Versus EPI Spatial Normalization Templates for fMRI Data Analyses. Hum Brain Mapp. 2017;38(11):5331–42.

5. Ding ZH, Huang YL, Bailey SK, Gao YR, Cutting LE, Rogers BP, et al. Detection of synchronous brain activity in white matter tracts at rest and under functional loading. Proceedings of the National Academy of Sciences of the United States of America. 2018;115(3):595–600.

6. Marussich L, Lu KH, Wen HG, Liu ZM. Mapping white-matter functional organization at rest and during naturalistic visual perception. NeuroImage. 2017;146:1128–41.

7. Huang Y, Yang Y, Hao L, Hu X, Wang P, Ding Z, et al. Detection of functional networks within white matter using independent component analysis. Neuroimage. 2020;222:117278.

8. Zhao YJ, Zhang FF, Zhang WJ, Chen LZ, Chen ZQ, Lui S, et al. Decoupling of Gray and White Matter Functional Networks in Medication-Naive Patients With Major Depressive Disorder. Journal of Magnetic Resonance Imaging. 2021;53(3):742–52.

9. Jiang Y, Song L, Li X, Zhang Y, Chen Y, Jiang S, et al. Dysfunctional white-matter networks in medicated and unmedicated benign epilepsy with centrotemporal spikes. Hum Brain Mapp. 2019;40(10):3113–24.

10. Chen X, Zhang H, Zhang L, Shen C, Lee SW, Shen D. Extraction of dynamic functional connectivity from brain grey matter and white matter for MCI classification. Hum Brain Mapp. 2017;38(10):5019–34.

11. Zhao J, Ding XT, Du YH, Wang XH, Men GZ. Functional connectivity between white matter and gray matter based on fMRI for Alzheimer’s disease classification. Brain Behav. 2019;9(10).

12. Gao Y, Sengupta A, Li M, Zu Z, Rogers BP, Anderson AW, et al. Functional connectivity of white matter as a biomarker of cognitive decline in Alzheimer’s disease. Plos One. 2020;15(10):e0240513.

13. Dai Z, Yan C, Li K, Wang Z, Wang J, Cao M, et al. Identifying and Mapping Connectivity Patterns of Brain Network Hubs in Alzheimer’s Disease. Cereb Cortex. 2015;25(10):3723–42.

14. Zhang H, Sachdev PS, Thalamuthu A, He Y, Xia M, Kochan NA, et al. The relationship between voxel-based metrics of resting state functional connectivity and cognitive performance in cognitively healthy elderly adults. Brain Imaging Behav. 2018;12(6):1742–58.

15. Tomasi D, Volkow ND. Functional connectivity density mapping. Proc Natl Acad Sci U S A. 2010;107(21):9885–90.

16. Buckner RL, Sepulcre J, Talukdar T, Krienen FM, Liu H, Hedden T, et al. Cortical hubs revealed 21 by intrinsic functional connectivity: mapping, assessment of stability, and relation to Alzheimer’s disease. J Neurosci. 2009;29(6):1860–73.

17. de Haan W, Mott K, van Straaten EC, Scheltens P, Stam CJ. Activity dependent degeneration explains hub vulnerability in Alzheimer’s disease. PLoS Comput Biol. 2012;8(8):e1002582.

18. Alexander-Bloch A, Raznahan A, Bullmore E, Giedd J. The convergence of maturational change and structural covariance in human cortical networks. J Neurosci. 2013;33(7):2889–99.

19. Marussich L, Lu KH, Wen H, Liu Z. Mapping white-matter functional organization at rest and during naturalistic visual perception. Neuroimage. 2017;146:1128–41.

20. Sepulcre J, Liu H, Talukdar T, Martincorena I, Yeo BT, Buckner RL. The organization of local and distant functional connectivity in the human brain. PLoS Comput Biol. 2010;6(6):e1000808.

21. Wu TL, Wang F, Li M, Schilling KG, Gao Y, Anderson AW, et al. Resting-state white matter-cortical connectivity in non-human primate brain. Neuroimage. 2019;184:45–55.

22. Bullmore E, Sporns O. The economy of brain network organization. Nat Rev Neurosci. 2012;13(5):336–49.

23. Liang X, Zou Q, He Y, Yang Y. Coupling of functional connectivity and regional cerebral blood flow reveals a physiological basis for network hubs of the human brain. Proc Natl Acad Sci U S A. 2013;110(5):1929–34.

24. Jack CR, Bernstein MA, Fox NC, Thompson P, Alexander G, Harvey D, et al. The Alzheimer’s Disease Neuroimaging Initiative (ADNI): MRI methods. J Magn Reson Imaging. 2008;27(4):685–91.

25. Weiner MW, Veitch DP, Aisen PS, Beckett LA, Cairns NJ, Green RC, et al. The Alzheimer’s Disease Neuroimaging Initiative 3: Continued innovation for clinical trial improvement. Alzheimers Dement. 2017;13(5):561–71.

26. Gu Y, Lin Y, Huang L, Ma J, Zhang J, Xiao Y, et al. Abnormal dynamic functional connectivity in Alzheimer’s disease. CNS neuroscience & therapeutics. 2020;26(9):962–71.

27. Mondragon JD, Maurits NM, De Deyn PP, Neur AD. Functional connectivity differences in Alzheimer’s disease and amnestic mild cognitive impairment associated with AT(N) classification and anosognosia. Neurobiol Aging. 2021;101:22–39.

28. Peer M, Nitzan M, Bick AS, Levin N, Arzyt S. Evidence for Functional Networks within the Human Brain’s White Matter. J Neurosci. 2017;37(27):6394–407.

29. Jiang Y, Luo C, Li X, Li Y, Yang H, Li J, et al. White-matter functional networks changes in patients with schizophrenia. NeuroImage. 2019;190:172–81.

30. Hojjati SH, Ebrahimzadeh A, Babajani-Feremi A, Initia ADN. Identification of the Early Stage of Alzheimer’s Disease Using Structural MRI and Resting-State fMRI. Frontiers in neurology. 2019;10.

31. Lee J, Ko W, Kang E, Suk HI, Initia ADN. A unified framework for personalized regions selection and functional relation modeling for early MCI identification. Neuroimage. 2021;236.

32. Olivito G, Serra L, Marra C, Di Domenico C, Caltagirone C, Toniolo S, et al. Cerebellar dentate nucleus functional connectivity with cerebral cortex in Alzheimer’s disease and memory: a seed-based approach. Neurobiol Aging. 2020;89:32–40.

33. de Reus MA, van den Heuvel MP. The parcellation-based connectome: limitations and extensions. Neuroimage. 2013;80:397–404. 22

34. Hayasaka S, Laurienti PJ. Comparison of characteristics between region-and voxel-based network analyses in resting-state fMRI data. Neuroimage. 2010;50(2):499–508.

35. Du HX, Liao XH, Lin QX, Li GS, Chi YZ, Liu X, et al. Test-retest reliability of graph metrics in high-resolution functional connectomics: a resting-state functional MRI study. CNS neuroscience & therapeutics. 2015;21(10):802–16.

36. Deng Y, Liu K, Shi L, Lei Y, Liang P, Li K, et al. Identifying the Alteration Patterns of Brain Functional Connectivity in Progressive Mild Cognitive Impairment Patients: A Longitudinal Whole-Brain Voxel-Wise Degree Analysis. Front Aging Neurosci. 2016;8:195.

37. Sheng J, Shen Y, Qin Y, Zhang L, Jiang B, Li Y, et al. Spatiotemporal, metabolic, and therapeutic characterization of altered functional connectivity in major depressive disorder. Hum Brain Mapp. 2018;39(5):1957–71.

38. Dai ZJ, Yan CG, Li KC, Wang ZQ, Wang JH, Cao M, et al. Identifying and Mapping Connectivity Patterns of Brain Network Hubs in Alzheimer’s Disease. Cereb Cortex. 2015;25(10):3723–42.

39. Chao-Gan Y, Yu-Feng Z. DPARSF: A MATLAB Toolbox for “Pipeline” Data Analysis of Resting-State fMRI. Frontiers in systems neuroscience. 2010;4:13.

40. Bartolomeo P, Thiebaut de Schotten M, Chica AB. Brain networks of visuospatial attention and their disruption in visual neglect. Front Hum Neurosci. 2012;6:110.

41. George K, J MD. Neuroanatomy, Thalamocortical Radiations. StatPearls. Treasure Island (FL) 2022.

42. Schulte T, Muller-Oehring EM. Contribution of callosal connections to the interhemispheric integration of visuomotor and cognitive processes. Neuropsychol Rev. 2010;20(2):174–90.

43. Yang L, Zhao C, Xiong Y, Zhong S, Wu D, Peng S, et al. Callosal Fiber Length Scales with Brain Size According to Functional Lateralization, Evolution, and Development. J Neurosci. 2022;42(17):3599–610.

44. Chan-Seng E, Moritz-Gasser S, Duffau H. Awake mapping for low-grade gliomas involving the left sagittal stratum: anatomofunctional and surgical considerations. J Neurosurg. 2014;120(5):1069–77.

45. Altieri R, Melcarne A, Junemann C, Zeppa P, Zenga F, Garbossa D, et al. Inferior Fronto-Occipital fascicle anatomy in brain tumor surgeries: From anatomy lab to surgical theater. J Clin Neurosci. 2019;68:290–4.

46. Ebeling U, Reulen HJ. Neurosurgical topography of the optic radiation in the temporal lobe. Acta Neurochir (Wien). 1988;92(1-4):29–36.

47. Mandonnet E, Martino J, Sarubbo S, Corrivetti F, Bouazza S, Bresson D, et al. Neuronavigated Fiber Dissection with Pial Preservation: Laboratory Model to Simulate Opercular Approaches to Insular Tumors. World Neurosurg. 2017;98:239–42.

48. Rayhan RU, Stevens BW, Timbol CR, Adewuyi O, Walitt B, VanMeter JW, et al. Increased brain white matter axial diffusivity associated with fatigue, pain and hyperalgesia in Gulf War illness. PLoS One. 2013;8(3):e58493.

49. Leunissen I, Coxon JP, Caeyenberghs K, Michiels K, Sunaert S, Swinnen SP. Task switching in traumatic brain injury relates to cortico-subcortical integrity. Hum Brain Mapp. 2014;35(5):2459–69. 23

50. Moeller K, Willmes K, Klein E. A review on functional and structural brain connectivity in numerical cognition. Front Hum Neurosci. 2015;9.

51. Radoeva PD, Jenkins GA, Schettini E, Gilbert AC, Barthelemy CM, DeYoung LLA, et al. White matter correlates of cognitive flexibility in youth with bipolar disorder and typically developing children and adolescents. Psychiatry research Neuroimaging. 2020;305:111169.

52. Sasson E, Doniger GM, Pasternak O, Tarrasch R, Assaf Y. Structural correlates of cognitive domains in normal aging with diffusion tensor imaging. Brain structure & function. 2012;217(2):503–15.

53. Dajani DR, Uddin LQ. Demystifying cognitive flexibility: Implications for clinical and developmental neuroscience. Trends in neurosciences. 2015;38(9):571–8.

54. Kim C, Cilles SE, Johnson NF, Gold BT. Domain general and domain preferential brain regions associated with different types of task switching: a meta-analysis. Human brain mapping. 2012;33(1):130–42.

55. Niendam TA, Laird AR, Ray KL, Dean YM, Glahn DC, Carter CS. Meta-analytic evidence for a superordinate cognitive control network subserving diverse executive functions. Cognitive, affective & behavioral neuroscience. 2012;12(2):241–68.

56. Yin RH, Tan L, Liu Y, Wang WY, Wang HF, Jiang T, et al. Multimodal Voxel-Based Meta-Analysis of White Matter Abnormalities in Alzheimer’s Disease. J Alzheimers Dis. 2015;47(2):495–507.

57. Ouyang X, Chen KW, Yao L, Wu X, Zhang JC, Li K, et al. Independent Component Analysis-Based Identification of Covariance Patterns of Microstructural White Matter Damage in Alzheimer’s Disease. Plos One. 2015;10(3).

58. Birdsill AC, Koscik RL, Jonaitis EM, Johnson SC, Okonkwo OC, Hermann BP, et al. Regional white matter hyperintensities: aging, Alzheimer’s disease risk, and cognitive function. Neurobiol Aging. 2014;35(4):769–76.

59. De Benedictis A, Duffau H, Paradiso B, Grandi E, Balbi S, Granieri E, et al. Anatomo-functional study of the temporo-parieto-occipital region: dissection, tractographic and brain mapping evidence from a neurosurgical perspective. J Anat. 2014;225(2):132–51.

60. Vannini P, Lehmann C, Dierks T, Jann K, Viitanen M, Wahlund LO, et al. Failure to modulate neural response to increased task demand in mild Alzheimer’s disease: fMRI study of visuospatial processing. Neurobiol Dis. 2008;31(3):287–97.

61. Prvulovic D, Hubl D, Sack AT, Melillo L, Maurer K, Frolich L, et al. Functional imaging of visuospatial processing in Alzheimer’s disease. Neuroimage. 2002;17(3):1403–14.

62. Tuch DS, Salat DH, Wisco JJ, Zaleta AK, Hevelone ND, Rosas HD. Choice reaction time performance correlates with diffusion anisotropy in white matter pathways supporting visuospatial attention. Proceedings of the National Academy of Sciences of the United States of America. 2005;102(34):12212–7.

63. Saur R, Milian M, Erb M, Eschweiler GW, Grodd W, Leyhe T. Cortical activation during clock reading as a quadratic function of dementia state. J Alzheimers Dis. 2010;22(1):267–84.

64. Ward A, Caro JJ, Kelley H, Eggleston A, Molloy W. Describing cognitive decline of patients at the mild or moderate stages of Alzheimer’s disease using the Standardized MMSE. Int Psychogeriatr. 2002;14(3):249–58. 24

65. Ferreira LK, Busatto GF. Resting-state functional connectivity in normal brain aging. Neuroscience and biobehavioral reviews. 2013;37(3):384–400.

66. Tomasi D, Volkow ND. Aging and functional brain networks. Mol Psychiatry. 2012;17(5):471, 549-58.

67. Reuter-Lorenz PA, Cappell KA. Neurocognitive aging and the compensation hypothesis. Curr Dir Psychol Sci. 2008;17(3):177–82.

68. Park DC, Reuter-Lorenz P. The Adaptive Brain: Aging and Neurocognitive Scaffolding. Annu Rev Psychol. 2009;60:173–96.

69. Qi Z, Wu X, Wang Z, Zhang N, Dong H, Yao L, et al. Impairment and compensation coexist in amnestic MCI default mode network. Neuroimage. 2010;50(1):48–55.

70. Zhang X, Guan Q, Li Y, Zhang J, Zhu W, Luo Y, et al. Aberrant Cross-Tissue Functional Connectivity in Alzheimer’s Disease: Static, Dynamic, and Directional Properties. J Alzheimers Dis. 2022;88(1):273–90.

71. Mori S, Oishi K, Jiang H, Jiang L, Li X, Akhter K, et al. Stereotaxic white matter atlas based on diffusion tensor imaging in an ICBM template. Neuroimage. 2008;40(2):570–82.

72. Kravitz DJ, Saleem KS, Baker CI, Mishkin M. A new neural framework for visuospatial processing. Nat Rev Neurosci. 2011;12(4):217–30.

73. Seeley WW, Menon V, Schatzberg AF, Keller J, Glover GH, Kenna H, et al. Dissociable intrinsic connectivity networks for salience processing and executive control. J Neurosci. 2007;27(9):2349–56.

74. Peters SK, Dunlop K, Downar J. Cortico-Striatal-Thalamic Loop Circuits of the Salience Network: A Central Pathway in Psychiatric Disease and Treatment. Front Syst Neurosci. 2016;10:104.

75. Di Carlo DT, Benedetto N, Duffau H, Cagnazzo F, Weiss A, Castagna M, et al. Microsurgical anatomy of the sagittal stratum. Acta neurochirurgica. 2019;161(11):2319–27.

76. Saper CB. The central autonomic nervous system: conscious visceral perception and autonomic pattern generation. Annu Rev Neurosci. 2002;25:433–69.

77. Seeley WW. The Salience Network: A Neural System for Perceiving and Responding to Homeostatic Demands. J Neurosci. 2019;39(50):9878–82.

78. Chand GB, Wu J, Hajjar I, Qiu D. Interactions of the Salience Network and Its Subsystems with the Default-Mode and the Central-Executive Networks in Normal Aging and Mild Cognitive Impairment. Brain Connect. 2017;7(7):401–12.

79. Fredericks CA, Sturm VE, Brown JA, Hua AY, Bilgel M, Wong DF, et al. Early affective changes and increased connectivity in preclinical Alzheimer’s disease. Alzheimers Dement (Amst). 2018;10:471–9.

80. Zaima N, Goto-Inoue N, Hayasaka T, Setou M. Application of imaging mass spectrometry for the analysis of Oryza sativa rice. Rapid communications in mass spectrometry: RCM. 2010;24(18):2723–9.

81. Liu Y, Yu C, Zhang X, Liu J, Duan Y, Alexander-Bloch AF, et al. Impaired long distance functional connectivity and weighted network architecture in Alzheimer’s disease. Cerebral cortex. 2014;24(6):1422–35.

